# A scalable platform for functional interrogation of peptide hormones in fish

**DOI:** 10.1101/2023.01.19.524675

**Authors:** Eitan Moses, Itamar Harel

## Abstract

Fish display a remarkable diversity of life-history traits, including body size, age at maturity, and longevity. Although pituitary hormones are conserved mediators of life-history transitions, regulatory networks are less understood in fish. However, the relatively long life-cycles and germline-dependent maturation of classical fish models are less compatible with rapid exploration of adult physiology, particularly in females. Here, we describe a high-throughput platform that combines, for the first time, loss- and gain-of-function of peptide hormones in a naturally short-lived fish. As a proof-of-principle, we first manipulate growth by mutating growth hormone (*gh1*) in the turquoise killifish (*N. furzeri*). Next, to rescue growth defects, we designed a vector in which hormones are tagged by a self-cleavable fluorescent reporter, and are ectopically expressed using intramuscular electroporation. A single injection of a *gh1-T2A-GFP* plasmid was sufficient to produce a stable expression of tag-free hormone and rescue growth phenotypes. This, in contrast to current practice for which multiple injections of recombinant hormones are required. We demonstrate the versatility of our platform by rescuing female sterility, which is induced by manipulating the follicle stimulating hormone (*fshb*). As killifish maturation is germline-independent, both sexes can be explored in genetic models with germline defects. Finally, we describe a doxycycline-inducible system for tunable expression control. Together, this platform significantly advances the state-of-the-art by allowing high-throughput functional dissection of distinct life-history strategies in fish. This method could be multiplexed to facilitate various applications, including optimizing commercially valuable traits in aquaculture, or screening pro-longevity hormonal interventions in aging.

## Introduction

Pituitary hormones are key regulators of vertebrate physiology, including growth (hypothalamic-pituitary-somatic axis), reproduction (hypothalamic-pituitary-gonadal axis), and metabolism (hypothalamic–pituitary–thyroid axis, **Figure 1a**). However, even though these hormones are largely conserved from fish to humans, regulatory networks are diverse and less understood in fish^1–4^. For example, while the loss of growth hormone (*gh1*) causes male infertility in zebrafish, a similar mutation in mice does not significantly affect reproduction^2,5,6^.

**Figure 1:**
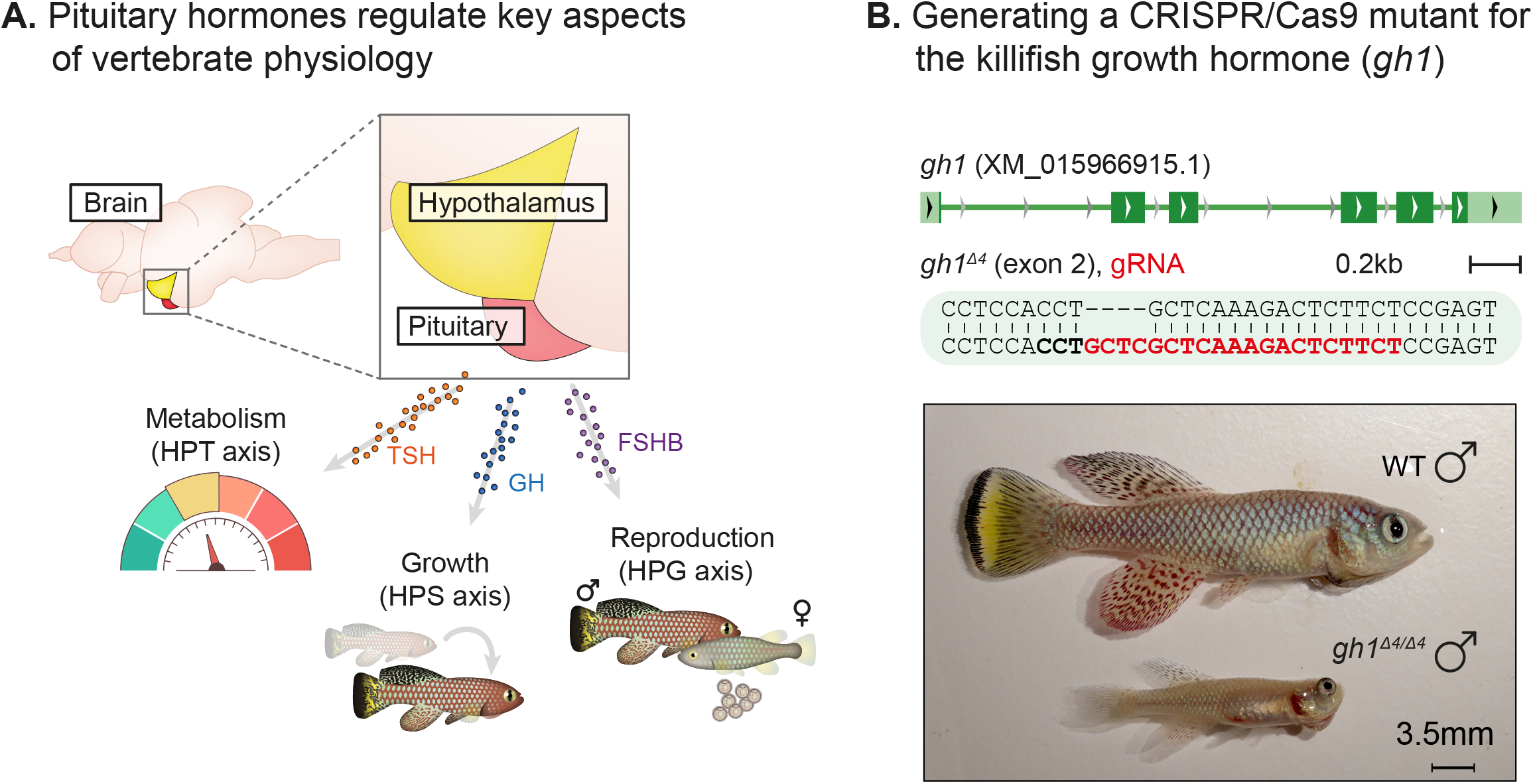
Perturbation of the killifish hypothalamic-pituitary-somatic axis. **(a)** Schematic illustration of the vertebrate hypothalamic-pituitary system, including members of the hypothalamic-pituitary-gonadal axis (HPG), the hypothalamic-pituitary-somatic axis (HPS), and hypothalamic-pituitary-thyroid axis (HPT). These hormones are released from the pituitary, and travel through the bloodstream to bind/activate their target receptors/organs. *fshb:* follicle stimulating hormone subunit beta; *gh1*: growth hormone; *tshb*: thyroid stimulating hormone subunit beta. **(b)** Generation of CRISPR mutants for *gh1*, with the guide RNA (gRNA) targets (red), protospacer adjacent motif (PAM, in bold), and indels (top). Bottom: Comparison of fish size between 8-week-old WT and *gh1*^*Δ4/Δ4*^ male fish.

Zebrafish (*Danio rerio*) and medaka (*Oryzias latipes*) are the most widely used genetic fish models. Interestingly, while these versatile models share a range of experimental advantages, they also exhibit several characteristics that are less compatible with high-throughput exploration of adult physiology. For example, both fish undergo relatively slow life cycles and sexual maturation (∼3-4 months), and depend on the presence of the germline to develop into females^7,8^. Accordingly, perturbations that affect the germline, such as mutation in the follicle stimulating hormone receptor gene (*fshr*), produce an all-male sex-reversal in both fish^7,8^ (while casing ovarian failure in humans^9^). Thus, as many peptide hormones also affect gonadal development, it might by challenging to explore both male and female physiology using these classical models.

Genetic perturbations of pituitary hormones in fish have been instrumental for understanding their function^1,2^. Yet, as these methods apply whole-body loss-of-function, interpretation could be biased from either stage-dependent hormonal requirements or potential compensatory mechanisms. Therefore, developing efficient rescue (or gain-of-function) of pituitary hormones is imperative for mechanistic understanding, and for determining specific temporal windows. Experimental approaches employed to date generally depend on laborious and repeated injections of pituitary extracts (which are non-specific), or require time-consuming purification of recombinant hormones^10,11^. Such issues significantly hamper the scalability of these experiments.

Here, we use the naturally short-lived turquoise killifish that exhibits one of the fastest recorded time to puberty among all vertebrate species (2-3 weeks, ∼6-fold faster compared to zebrafish and medaka). An underexplored advantage of the killifish model is that sexual maturation is germline-independent^12^. Thus, allowing us to explore both males and females in a wide range of hormonal manipulations that can affect germline development. Mutating the killifish *gh1* significantly inhibits somatic growth and delays maturity. We then developed a vector system in which a desired hormone is tagged using a self-cleavable fluorescent reporter, and ectopically expressed *in-vivo* using intramuscular electroporation. Following electroporation, *gh1* was stably expressed and partially rescued growth defects. In addition, we demonstrate the versatility of this approach in other pituitary hormones, and demonstrate the feasibility of dose-dependent and doxycycline-inducible systems for accurate temporal expression patterns. Our platform therefore represents a simple and robust method for loss/gain-of-function of circulating factors in fish. Importantly, this strategy can be multiplexed, is effective after a single injection, and can be readily adapted for other fish species.

## Results

### Generation of a growth hormone CRISPR mutant in killifish

The hypothalamic–pituitary–somatic (HPS) axis includes the secretion of growth hormone (GH) from the pituitary gland, and the consequent stimulation of insulin-like growth factor 1 (IGF-1) production in peripheral tissues (primarily the liver). We applied our recently developed CRISPR/Cas9 genome editing protocols^12–15^ to perturb the HPS axis in the turquoise killifish by targeting exon 2 of the killifish *gh1* (**Figure 1b**, top). Crossing F0 founders with wild-type fish allowed us to identify a 4bp deletion in F1 fish that could cause a frame-shift. The *gh1*^*Δ4/+*^ heterozygous mutants were outcrossed for several generations to reduce the chances of off target alterations. The *gh1*^*Δ4/+*^ mutants were viable and fecund, and were used to generate homozygous *gh1*^*Δ4/Δ4*^ mutants for phenotypic analysis.

### Mutating the killifish growth hormone delays somatic growth and maturity

Both male and female *gh1*^*Δ4/Δ4*^ homozygous mutants were strikingly smaller than control fish (**Figures 1b, S1a**). The proportion of *gh1*^*Δ4/Δ4*^ mutants, out of the total number of genotyped individuals, was roughly half of the expected Mendelian distribution (p= 2×10^−5^ in males, and 9×10^−5^ in females, by Chi square, **Figure S1b**). Most of this reduction could be attributed to their significantly smaller size, since the smaller fish were more easily outcompeted in the communal tanks (or escaped) until they were individually housed following genotyping at 4 weeks of age. Interestingly, while both male and female mutants were significantly smaller, they were still fertile (**Figure 2a, b**). However, their slow growth was coupled with a roughly ∼2-fold delay in the onset of egg lay, male pigmentation, and a proportional delay in peak fertility (**Figures 1b, 2b**). As expected, the total number of eggs in each egg lay was also much smaller in the mutant couples, and is probably limited by the smaller female body size (**Figure 2b**).

**Figure 2:**
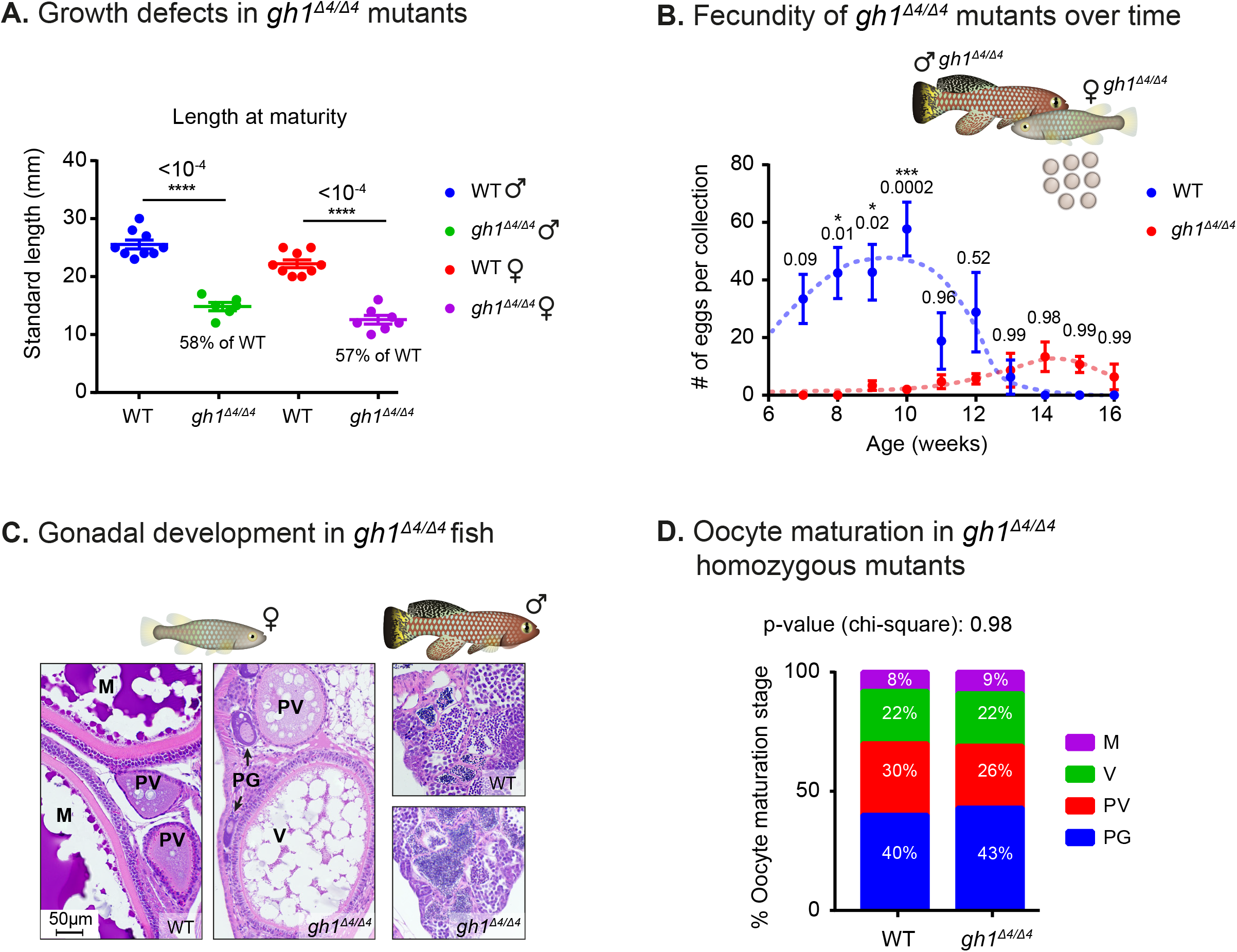
Phenotypical analysis of growth hormone mutants. **(a)** Quantification of somatic growth (standard length) of 8-week-old WT or *gh1*^*Δ4/Δ4*^ mutants: males (left) and females (right), n ≥ 6 individuals from each experimental group. Error bars show mean ± SEM. Significance was calculated using one-way ANOVA with a Sidak post-hoc comparing the male and female mutants to the respective WT. Exact p-values are indicated. The relative size of the mutant fish, when compared to the corresponding controls, is indicated as %. **(b)** Quantification of reproductive output in *gh1*^*Δ4/Δ4*^ mutant pairs. Each dot represents the mean number of eggs of the indicated genotypes, per week of egg collection. There were 3 independent mutant pairs and 7 WT pairs. Error bars show mean ± SEM. Significance was calculated using repeated measures two-way ANOVA with a Sidak post-hoc compared to the WT. Exact p-values are indicated. **(c)** Representative histological sections, depicting ovaries and testis of the indicated genotypes. n ≥ 4 individuals (two-month-old), from each genotype. Scale bar: 50 µm. **(d)** Distribution of oocyte development stages. Data are presented as the proportion of each developmental stage of the indicated genotypes. n ≥ 4 individuals for each experimental group. Significance was measured by χ^2^ test with the WT value as the expected model and FDR correction. Percentages and exact p-values are indicated. PG: primary growth; PV: pre-vitellogenic; V: vitellogenic; M: Mature.

As the next stage, we stained 2-month old fish gonads with Haematoxylin & Eosin (H&E). We then examined tissue sections for changes in germ cell development caused by GH deficiency (**Figure 2c**), and quantified the proportion of gonadal germ cells at each developmental stage in both females and males (**Figures 2d, S2a**). Interestingly, the results revealed that once the mutants reached maturity, oocyte and sperm development are generally unaffected by the deficiency of growth hormone. Specifically, oogenesis was comparable in mutant fish and sibling controls (**Figure 2d**), as was spermatogenesis (**Figure S2a**). Taken together, our findings demonstrate that GH deficiency delays both somatic growth and maturity.

### Developing an efficient and scalable method to rescue hormonal perturbations

In order to rescue GH deficiency, we developed a gain-of-function approach to ectopically express a gene of interest by optimizing intramuscular electroporation in killifish (**Figure 3a**, see **Methods**)^16–19^. *Gh1* from turquoise killifish cDNA was cloned upstream to GFP, and separated by the T2A self-cleaving peptide^20^ (**Figure 3a**, top). We injected the construct (∼3µg, in 3-5µl) into the muscle of WT fish, performed electroporation, and imaged GFP expression 72h post electroporation (**Figure 3a**, bottom). This approach enables faithful visualization of the electroporation efficiency, which serves as a proxy for hormone expression level. Importantly, since GH and GFP are co-translated, they are predicted to maintain a 1:1 ratio. The GFP tag is removed after the self-cleavage so that hormonal secretion and function is expected to be unaffected. Importantly, we were able to detect GH expression in GFP-positive muscle fibers by immunofluorescence (**Figure S3a**).

**Figure 3:**
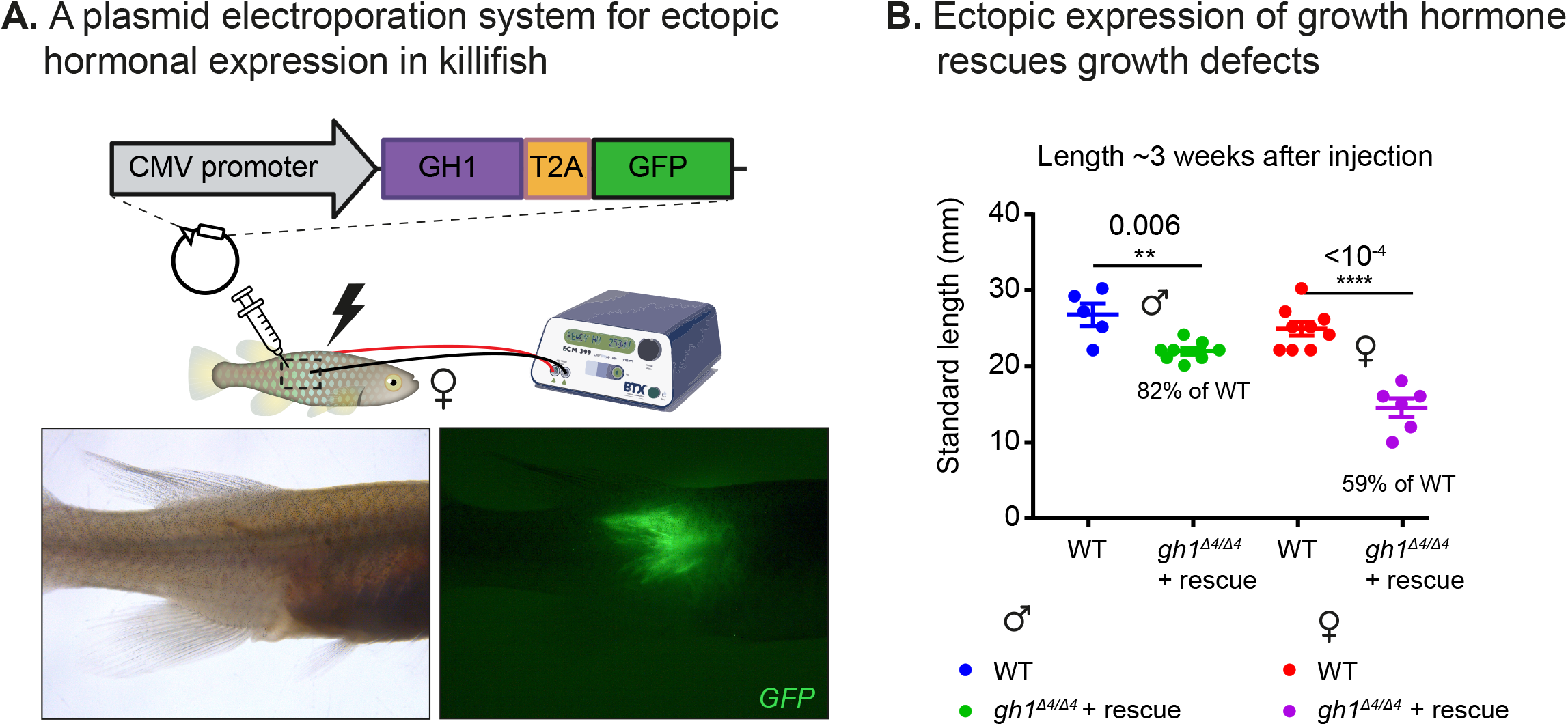
Phenotypic rescue of growth hormone deficiency by ectopic expression. **(a)** Schematic illustration of the gain-of-function plasmid, and intramuscular ectopic plasmid electroporation (top). GFP is visible following the electroporation of the GH:GFP plasmid (bottom). **(b)** Quantification of somatic growth (standard length) of 12-week-old *gh1*^*Δ4/Δ4*^ mutants following electroporation of a *gh1-T2A-GFP* plasmid as compared to WT controls: males (left) and females (right), n ≥ 5 individuals from each experimental group. Error bars show mean ± SEM. Significance was calculated using one-way ANOVA with a Sidak post-hoc comparing the male and female mutants to the respective WT animals. Exact p-values are indicated. The relative size of the mutant fish, when compared to the corresponding controls, is indicated as %.

Next, we injected the same construct into the muscle of 8-week-old male and female *gh1*^*Δ4/Δ4*^ fish, performed electroporation, and monitored their somatic growth. An increase in size was observed 3-4 weeks following electroporation, particularly in males (**Figure 3b**). As measured by the relative size of electroporated mutants compared to WT controls, the electroporated male mutants were 82% of size of age-matched controls (**Figure 3b**), compared to a value of only 58% for nonelectroporated mutant fish (**Figure 2a**). These results demonstrate that muscle electroporation can substantially rescue the hormonal deficiency phenotype.

### Reversible perturbation of the reproductive axis

The versatility of this approach was demonstrated in two other systems: manipulating killifish reproduction by mutating the follicle stimulating hormone (*fshb*), an integral part of the HPG axis (**Figures 4, S4a**) and affecting the HPT axis by mutating the thyroid stimulating hormone (*tshb*, **Figure S4a**). To evaluate the role of FSHB on killifish reproduction, we imaged haematoxylin and eosin (H&E) stained tissue sections from homozygous *fshb*^*in1/in1*^and WT females (n > 4 for each experimental condition). Our findings demonstrated a striking reduction in mature oocytes, which were essentially completely missing in homozygous *fshb*^*in1/in1*^ females (**Figure 4a**). Consequently, mating of *fshb*^*in1/in1*^ females with WT males confirmed that mutant females are generally infertile (**Figure 4b**).

**Figure 4:**
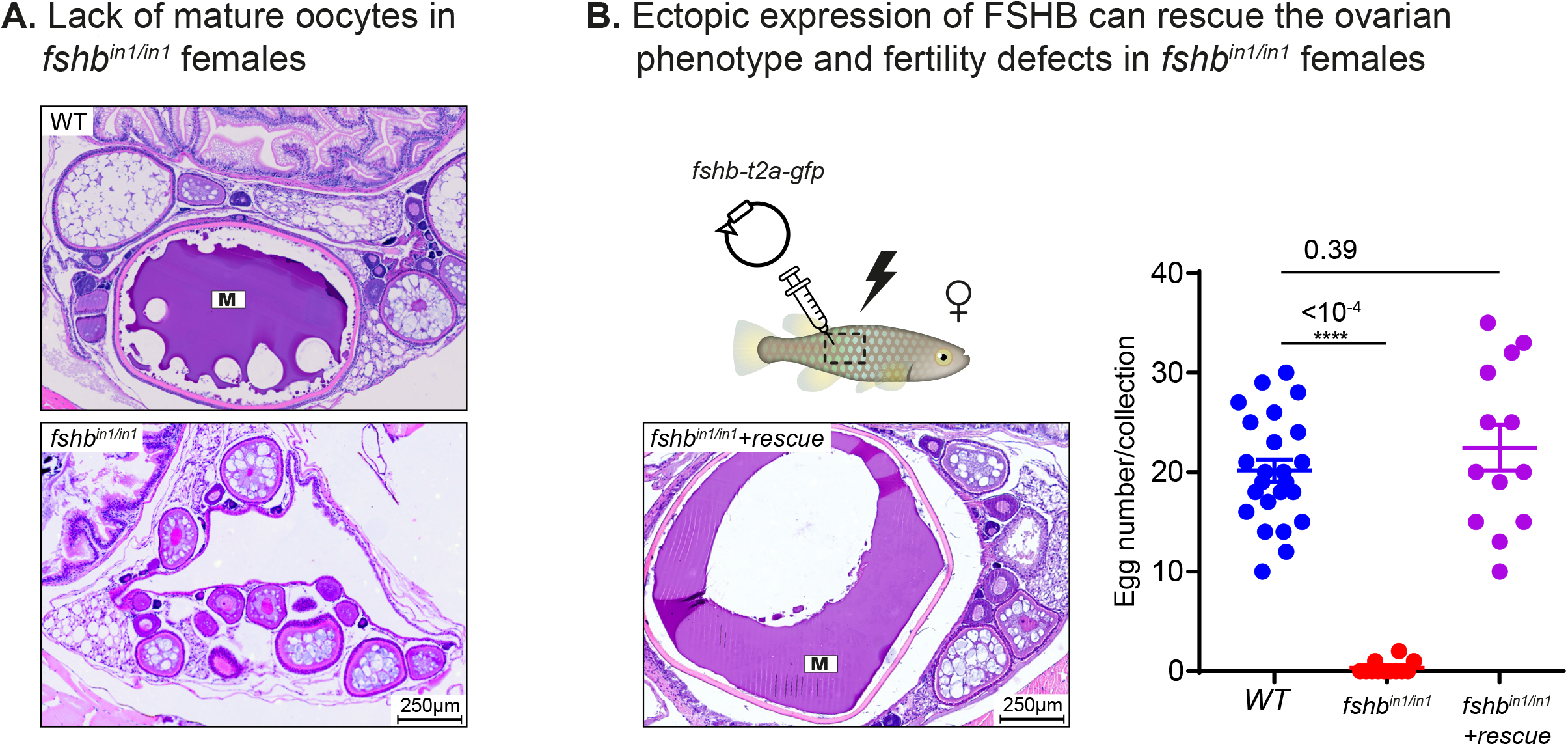
Reversible perturbation of the killifish reproductive axis. **(a)** Representative histological sections, depicting one-month-old ovaries of the indicated genotypes. n ≥ 4 individuals from each genotype. Scale bar: 250 µm. **(b)** Left: representative histological sections of ovaries in *fshb*^*in1/in1*^mutant females following electroporation of *fshb*-T2A-GFP. Representative of n ≥ 3 individuals. Right: quantification of female fertility. Each dot represents the number of eggs per breeding pair of the indicated experimental conditions, per week of egg collection. The data are from at least 3 independent pairs and 4 independent collections. Error bars show mean ± SEM with individual points. Significance was calculated using one-way ANOVA with a Tukey post-hoc and exact p-value is indicated.

While *fshb* killifish mutants displayed predominantly gonadal-related phenotypes (causing an arrest in female germ cell maturation), no significant effect on growth has been seen. In contrast, disrupted thyroid hormone signaling is expected to produce both reproductive defects (lack of mature oocytes) as well as growth defects in both mice^21^ and zebrafish^22^. Accordingly, in our system, we observed growth malformations, particularly in overall fish shape, in pigmentation, as well as defects in oocyte maturation in the homozygous *tshb*^*Δ10/Δ10*^ killifish mutants (**Figure S4b**).

Next, we selected the *fshb*^*in1/in1*^ to attempt and rescue the reproductive axis, as restoration of female infertility will provide a simple readout for the efficiency and robustness of our electroporation protocol. We first cloned *fshb* from turquoise killifish cDNA to generate a CMV: *fshb-T2A-GFP* plasmid. Intramuscular electroporation of the construct completely restored the fertility of *fshb*^*in1/in1*^ females. This fertility persisted for at least one month (**Figure 4b**), which suggests that the plasmid is stably expressed for several weeks. Detailed characterization of the *fshb*^*in1/in1*^ mutant fish and other pituitary hormones, and specifically the reproductive defects and their rescue, will be described elsewhere (E.M. and I.H., unpublished).

### Developing a tunable expression system that is compatible with pulsatile release

In humans, regulation of puberty and mensural cycles depends on precise temporal changes of hormone levels^23^. For example, in prepubescent girls, gonadotropin-releasing hormone (GnRH) is required to stimulate the release of LH and FSH. Ultimately, LH peak amplitude increases about 10-fold, whereas FSH pulse amplitude only doubles^23^ (see a schematic diagram in **Figure 5a**, top). After puberty, a particular sequence is also critical for ovulation (**Figure 5a**, bottom). Thus, to develop a tight control of expression levels in fish we tested different plasmid concentration. Specifically, we used the electroporation method described above, with total plasmid amount per fish ranging by 125-fold (3µg - 24ng). Imaging these fish 3 days after injection revealed a dose-dependent GFP expression (**Figure 5b**).

**Figure 5:**
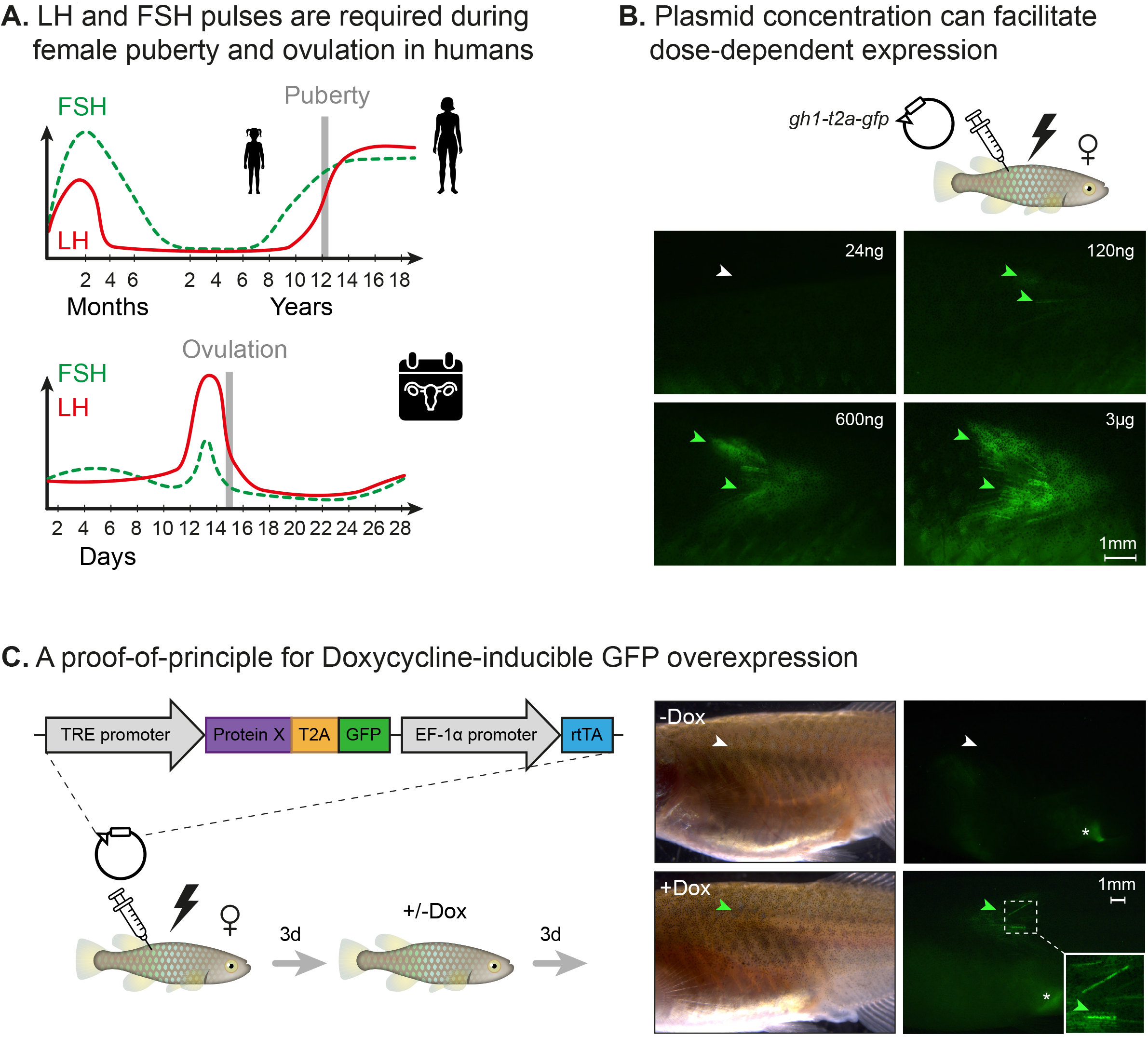
Dose dependent and inducible expression systems. **(a)** A schematic illustration depicting complex expression levels of LH (red line) and FSH (green line) during puberty (top) and ovulation (bottom) in human females. **(b)** Electroporation of the indicated plasmid concentrations. GFP signal (green arrowheads) and lack of signal (white arrowheads) are shown. n≥3 for each experimental condition. Scale bar = 1mm. **(c)** Left: schematic illustration showing electroporation of a plasmid with a doxycycline-inducible promoter, coding for a desired protein (ProteinX) and tagged with a T2A GFP. Right: following doxycycline treatment, a GFP signal is observed (green arrowheads). In control fish (-Dox), there is no detectable signal (white arrowheads). n≥3 for each experimental condition. Scale bar = 1mm. autofluorescence is marked by an asterisk.

While these experiments provide the means to control expression levels, some expression patters require delayed activation or pulsations (**Figure 5a**). Thus, we explored the Doxycycline inducible overexpression system (**Figure 5c**, left). This system includes a doxycycline-dependent transcriptional activator protein (rtTA) and a tet operator (tetO) containing promoter (TRE promoter)^24^. Only in the presence of doxycycline, rtTA can bind to the tetO promoter and activate gene expression. Specifically, we injected 3µg of a plasmid with a doxycycline-dependent GFP expression. Three days post electroporation, fish were exposed to doxycycline for 72h, and imaged. Our data indicated that this approach could be successfully applied for producing time- and dose-dependent hormone expression patterns (**Figure 5c**, right).

## Discussion

Here, using the turquoise killifish, we develop a platform to rapidly and reversibly manipulate life history traits in fish. The capabilities of the system are demonstrated by genetic perturbation of pituitary hormones, which are master regulators of vertebrate physiology. By mutating *gh1* and *fshb* we efficiently alter somatic growth and reproduction, respectively. We then develop a vector system that restores hormone function by ectopically expressing the missing hormones using intramuscular electroporation. Importantly, the naturally compressed life cycle of the killifish model significantly shortens experimental timescales, and allows for rapid physiological readouts of both loss- and gain-of function. Together, this method is relatively high-throughput, and thus facilitates large-scale interrogation of peptide hormones in fish.

Manipulation of the killifish GH and TSH alters both somatic growth and reproduction (**Figure 2a, b**). This apparent co-regulation is widely observed, and a number of evolutionary theories predict functional trade-offs between life-history traits^25–28^. For example, pituitary hormones link somatic growth with organismal lifespan in the long-lived Ames and Snell dwarf mice, which suffer from a deficiency of pituitary growth hormone (among other hormones)^29–31^. Similarly, mice with a mutated growth hormone receptor are long-lived^32^, and in humans, longer lives and cancer protection are observed in Laron Syndrome patients (dwarfism due to a growth hormone receptor mutation)^33^. These complex relationships raise interesting predictions. For example, can a specific delay in maturity drive proportional changes in vertebrate lifespan? Better understanding of these mechanisms will allow us to uncouple linked traits, such as somatic growth and reproduction (**Figure 6**).

**Figure 6:**
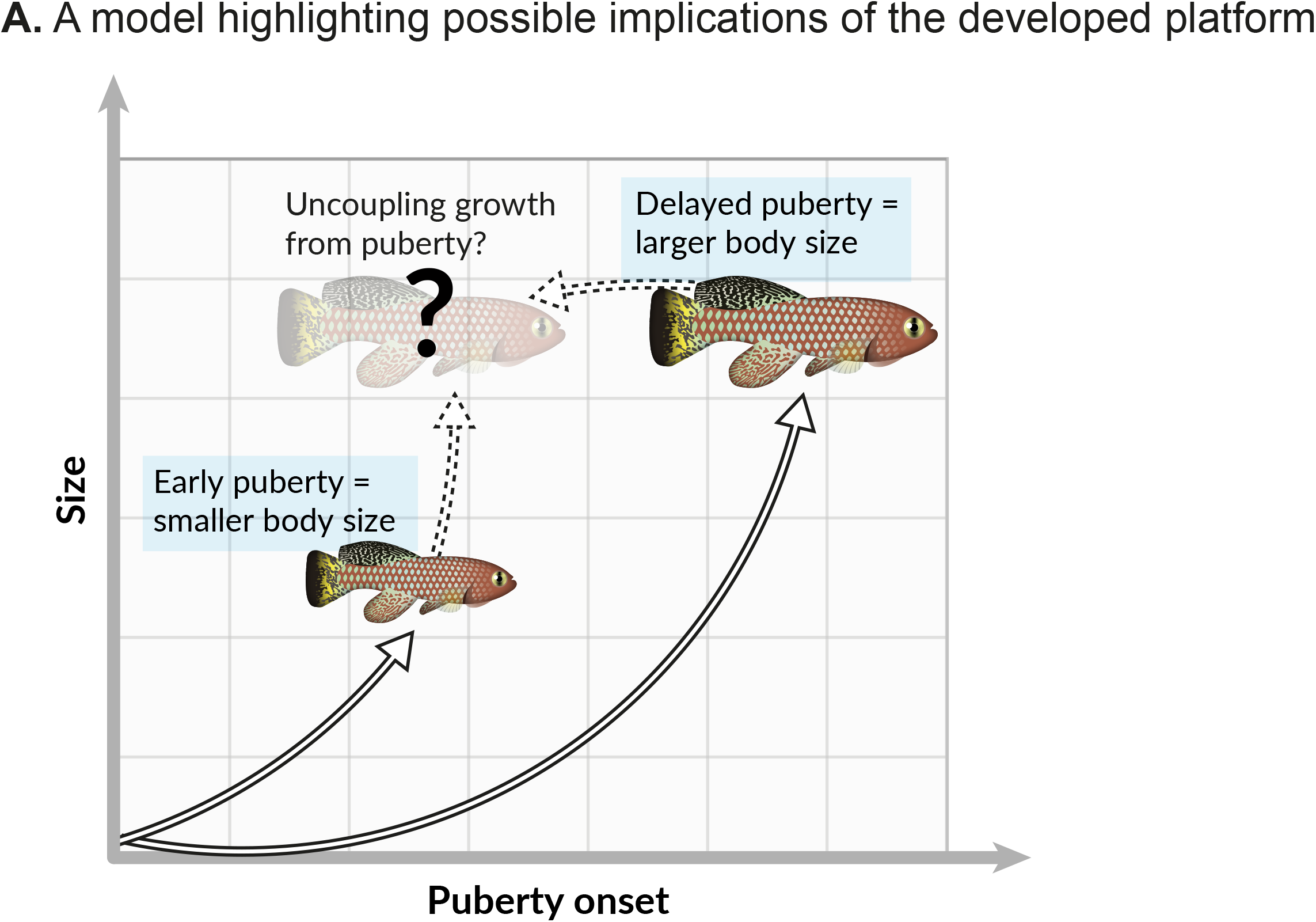
Possible implications of the developed platform. (**a**) Onset of maturity can negatively affect growth. Identifying the molecular mechanisms that regulate these seemingly opposing traits, and potentially uncoupling them, holds great promise for basic research and aquaculture.

So far, we and others have primarily perturbed a single hormone at a time. However, in reality, multiple systemic signals are integrated to determine the duration and onset of life-history traits. Our method is compatible with multiplexing, and could allow the examination of combinatorial effects of several hormones. Similarly, to explore requirements within specific temporal windows, the dose-dependent and doxycycline-inducible system could be used for a sequential and reversible gain-of-function.

Ultimately, these tools could facilitate the optimization of commercially valuable traits in aquaculture^11^. So far, the majority of fish species used in aquaculture are wild-caught, or have undergone only initial steps of selective breeding. Additionally, in some species, current manipulations are extremely time consuming (e.g. some eel species require ∼16 weekly injections of pituitary extracts to induce maturation^34^). Thus, plasmid electroporation, which does not modify the host genome, could replace traditional methods of hormone administration to manipulate many traits, including reproductive cycle, growth, and diseases resistance.

We have recently demonstrated that the killifish model could be used to identify novel regulators of aging, which involve systemic modulation of metabolism^14^. Therefore, beyond our current application, it would be exciting to expand this platform to investigate systemic factors as modifiers of health and disease. For example, recent studies have demonstrated that circulating factors in young plasma can rejuvenate old mice^35–44^. So far, only a handful of these factors have been identified, including VEGF^44^ (Vascular Endothelial Growth Factor), and GDF11^41^ (Growth Differentiation Factor 11). Thus, many possible candidates that arise from age-dependent changes in the plasma proteome remain to be explored^45^. In conclusion, our platform significantly advances the state-of the-art by providing the tools to functionally dissect the mechanisms that regulate vertebrate life history at an unprecedented resolution and speed.

## Figure legends

**Figure S1:**
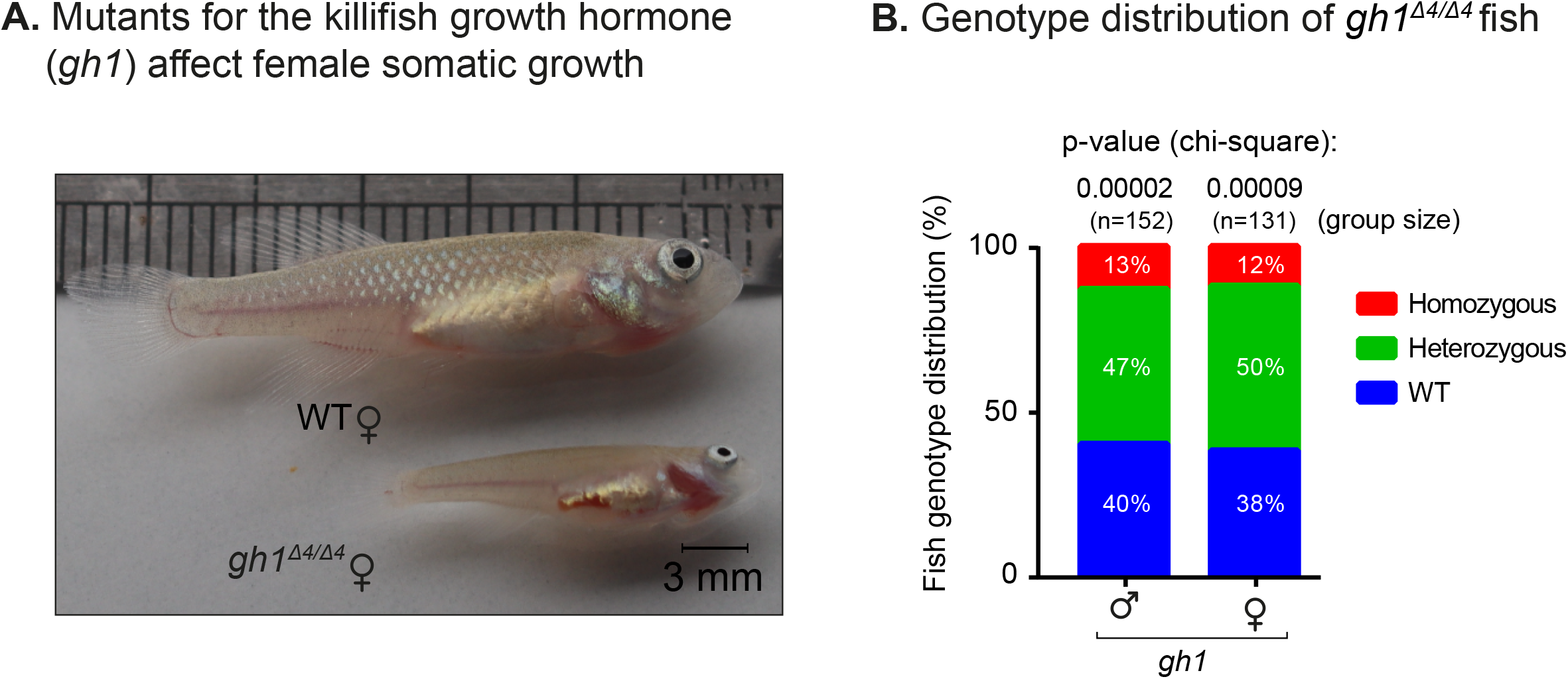
physiological effects of growth hormone deficiency. **(a)** Comparison of fish size between 6-week-old WT and *gh1*^*Δ4/Δ4*^ female fish. **(b)** Distribution of genotype progeny from heterozygous *gh1*^*Δ4/+*^ pairs (stratified by sex). n > 130 individuals, per sex. Percentages are indicated, and significance was measured by χ^2^ test with Mendelian proportions (25:50:25) as the expected model and FDR correction.

**Figure S2:**
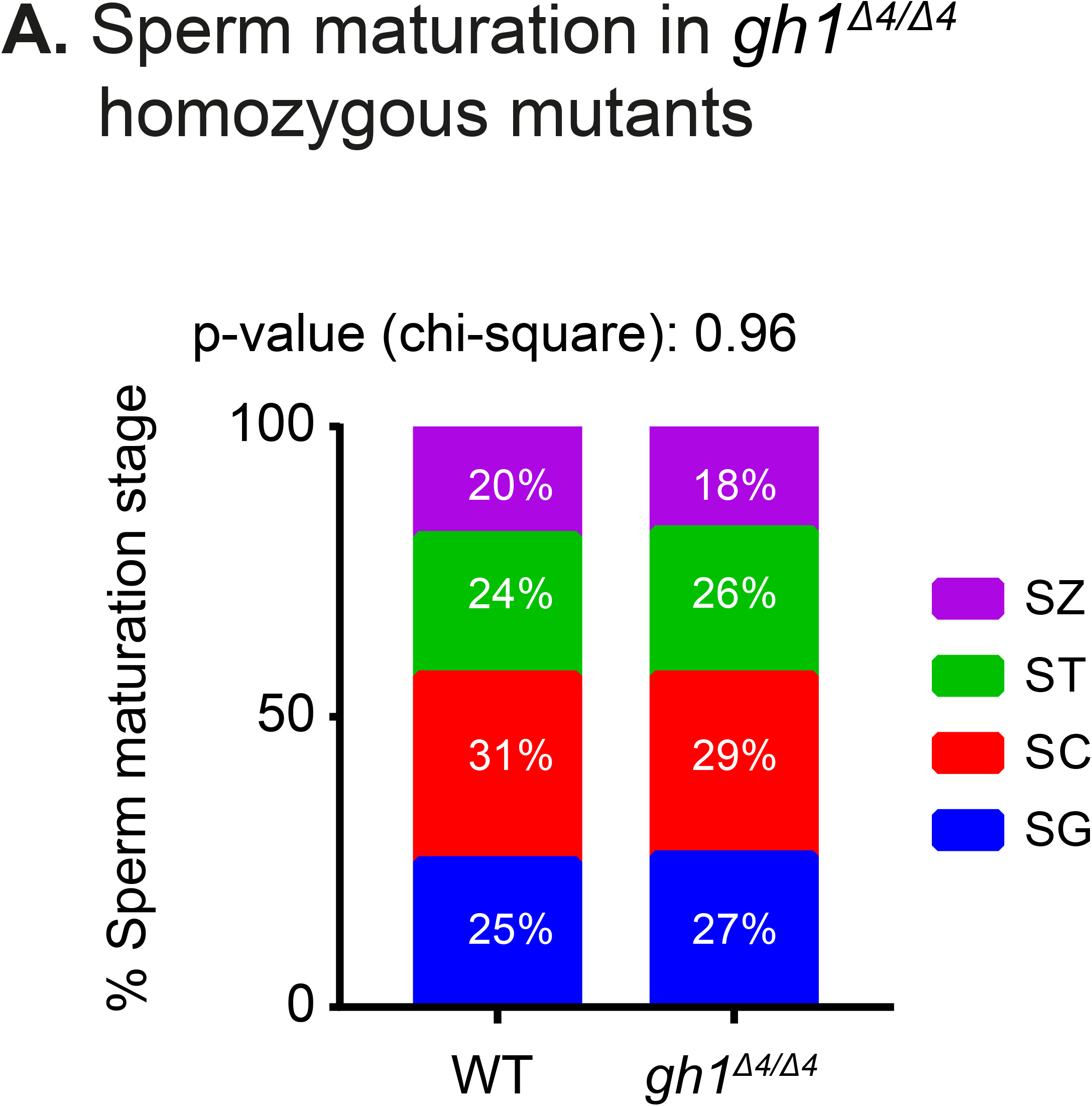
Sperm maturation in growth hormone mutants. (**a**) Quantification of sperm maturation, examples in (Figure 2c). Data are presented as the proportion of each developmental stage of the indicated genotypes. n ≥ 4 individuals for each experimental group. Significance was measured by χ^2^ test with the WT value as the expected model and FDR correction. Percentages and exact p-values are indicated. Sperm developmental stages according to^102^: SG: spermatogonia; SC: spermatocytes; ST: spermatids; SZ: spermatozoa.

**Figure S3:**
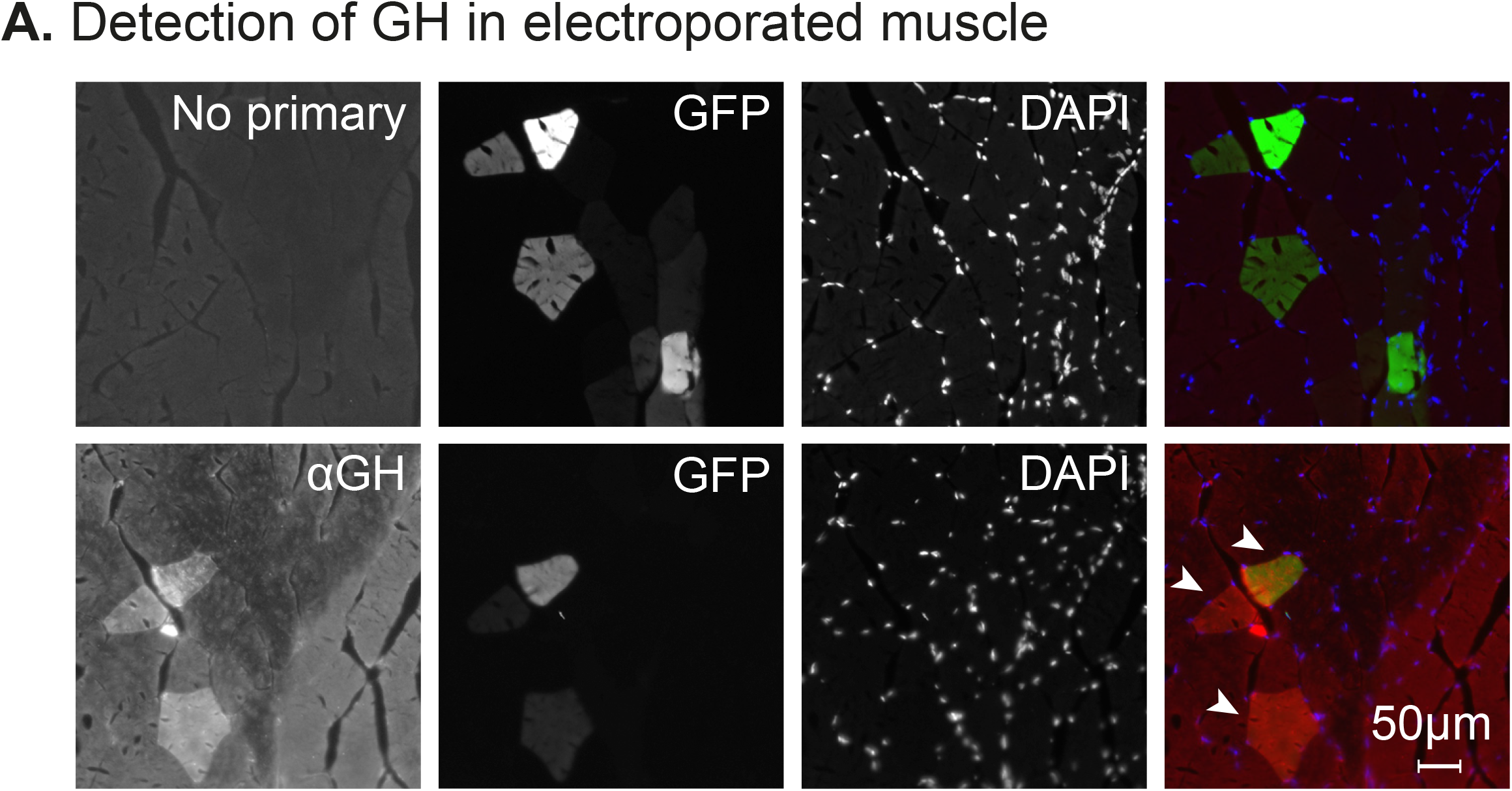
Design of growth hormone expressing plasmid. (**a**) Immunofluorescence for GH expression (red) on muscle cryosections of electroporated fish. GFP expression is shown in green, and nuclear staining (DAPI) in blue. Representative of 3 fish from each experimental group. Scale bar: 50µm.

**Figure S4:**
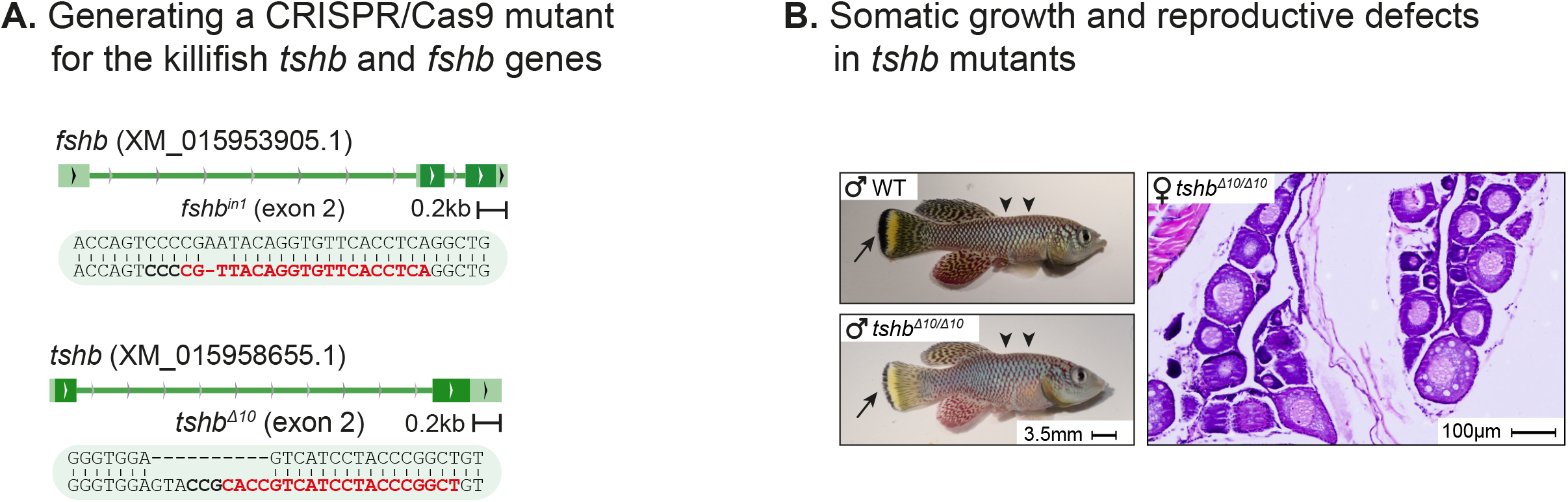
Generation of *fshb* and *tshb* mutants in killifish: **(a)** Generation of CRISPR mutants for *fshb* and *tshb*, depicting the guide RNA (gRNA) targets (red), protospacer adjacent motif (PAM, in bold), and indels (top). Scale bar: 3.5 mm. **(b)** Left: comparison of two-month-old WT and *tshb*^*Δ10/Δ10*^ male fish. Black arrows highlight tail melanocytes, while arrowheads indicated alterations in body shape. Right: representative histological sections demonstrating that one-month-old *tshb*^*Δ10/Δ10*^ ovaries lack mature oocytes. Scale bar: 100 µm.

**Table S1.**
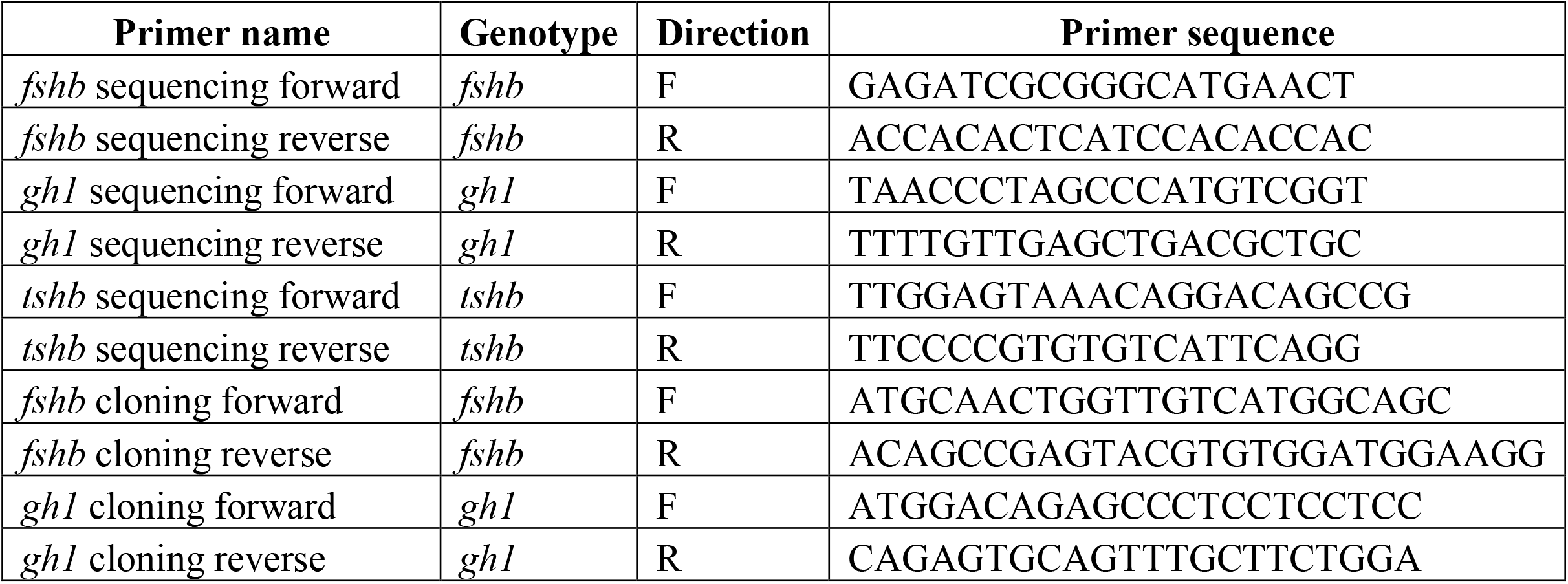

## Acknowledgements

We thank the Harel lab for a stimulating discussion and feedback on the manuscript. We thank Ariel Velan, Ella Yanay, and Ashayma Abu-tair for help with killifish maintenance and Adi Oron-Gottesman for her help with cloning. Supported by the Zuckerman Program (I.H.), ISF 2178/19 (I.H.), Israeli Ministry of Science 3-17631 (I.H.), 3-16872 (I.H.), the Moore Foundation GBMF9341 (I.H.), BSF-NSF 2020611 (I.H.), the Israeli Ministry of Agriculture 12-16-0010 (I.H) and the Levi Eshkol scholarship of the Israeli Ministry of Science (E.M).

## Author Contributions

E.M. and I.H. designed the study. E.M. performed experiments and generated the *gh1, fshb* and *tshb* mutant killifish lines. E.M. and I.H. wrote the manuscript.

## MATERIALS AND METHODS

### MATERIALS AVAILABILITY

All plasmids and corresponding annotated maps are available via Addgene (# 194356, #194883, #196331). All fish lines are available upon request.

## EXPERIMENTAL MODEL AND SUBJECT DETAILS

### Data analysis

No data was excluded during the analysis, as no significant outliers or classical exclusion criteria (e.g. unnatural death) occurred during experimentation. We assumed normality for all data, as commonly applied for physiological traits (such as size, fertility etc.). Therefore, a Student’s t-test could be used for comparing 2 groups, and ANOVA for more than 2 groups. In case of repeated measures over time, we used a 2-way repeated measures ANOVA test with time as one variable. To calculate proportions within a group we used a χ^2^ test. For biological replicates, we used parallel measurements of individual fish that captures random variation. Technical replicates were considered when the same experiment was conducted several times, such as several egg collections from the mating pair. Power analysis was performed for growth measurements, predicting a reduction of 50% in size with an alpha of 0.05 and power of 80%:

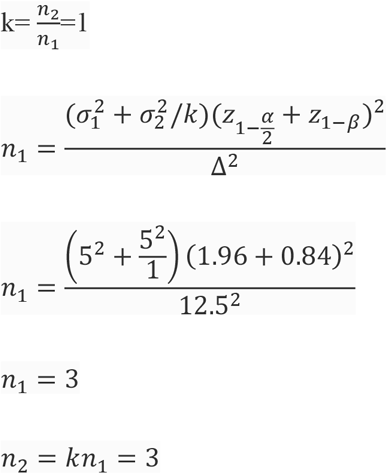

### African turquoise killifish strain, husbandry, and maintenance

The African turquoise killifish (GRZ strain) was housed as previously described^13,15^. Fish were grown at the Hebrew University of Jerusalem (Aquazone ltd, Israel) in a central filtration recirculating system at 28°C, with a 12 h light/dark cycle. Fish were fed with live Artemia until the age of 2 weeks (#109448, Primo), and starting week 3, fish were fed three times a day on weekdays (and once a day on weekends), with GEMMA Micro 500 Fish Diet (Skretting Zebrafish, USA), supplemented with Artemia once a day. All genetic models (described below) were maintained as heterozygous and propagated by crossing with wild-type fish. All turquoise killifish care and uses were approved by the Subcommittee on Research Animal Care at the Hebrew University of Jerusalem (IACUC protocols #NS-18-15397-2 and #NS-22-16915-3).

### CRISPR/Cas9 target prediction and gRNA synthesis

CRISPR/Cas9 genome-editing protocols were performed as described previously^13^. Briefly, evolutionary conserved regions upstream of functional or active protein domains were selected for targeting the selected genes. gRNA target sites were identified using CHOPCHOP (https://chopchop.rc.fas.harvard.edu/)^46^, and are shown below. PAM sites are shown in bold, and when needed, the first base-pair was changed to a ‘G’ to comply with the T7 promoter.

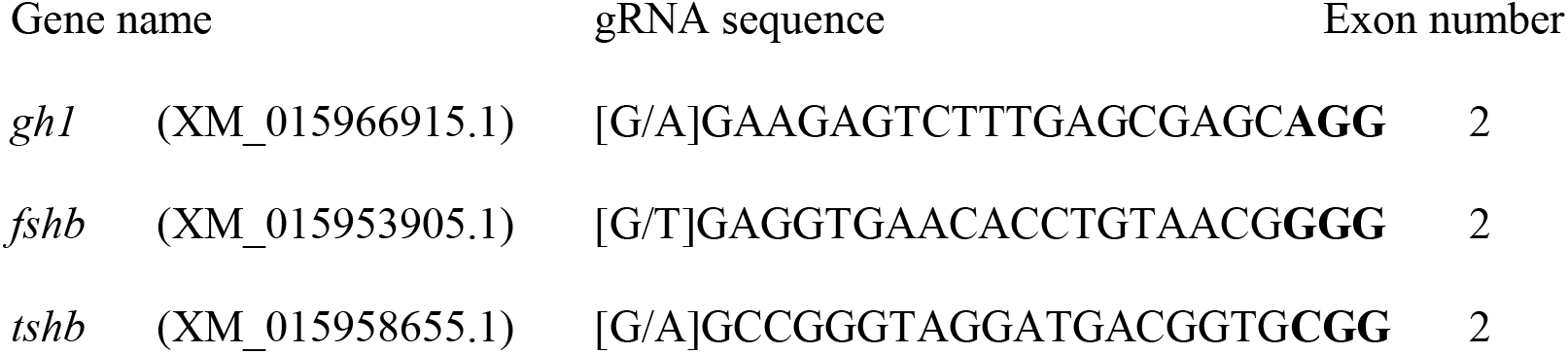

Design of variable oligonucleotides, and hybridization with a universal reverse oligonucleotide was performed according to^13^, and the resulting products were used as a template for *in vitro* transcription. gRNAs were *in vitro* transcribed and purified using a quarter reaction of TranscriptAid T7 High Yield Transcription Kit (Thermo Scientific #K0441), according to the manufacturer’s protocol.

### Production of Cas9 mRNA

Experiments were performed as described previously^12,13^. The pCS2-nCas9n expression vector was used to produce Cas9 mRNA (Addgene, #47929)^47^. Capped and polyadenylated Cas9 mRNA was *in vitro* transcribed and purified using the mMESSAGE mMACHINE SP6 ULTRA (ThermoFisher # AM1340).

### Microinjection of turquoise killifish embryos and generation of mutant fish using CRISPR/Cas9

Microinjection of turquoise killifish embryos was performed according to^13^. Briefly, nCas9n-encoding mRNA (300 ng/μL) and gRNA (30 ng/μL) were mixed with phenol-red (P0290, Sigma-Aldrich) and co-injected into one-cell stage fish embryos. Sanger DNA sequencing was used for detecting successful germline transmission on F1 embryos. Fish with desired alleles were outcrossed further to minimize potential off-target effects, and maintained as stable lines by genotyping using the KASP genotyping platform (Biosearch Technologies) with custom made primers. All primers used in generating the mutations can be found in Table S1.

## METHOD DETAILS

### Growth measurements

Both sexes were measured by imaging at the indicated timepoints with a Canon Digital camera EOS 250D, prime lens Canon EF 40 mm f/2.8 STM. To limit vertical movement during imaging, fish were placed in a tank with 3 cm deep water, and images were taken from the top using fixed lighting and height. A ruler was included in each image to provide an accurate scale. Body standard length was measured from the tip of the snout to the posterior end of the last vertebra (excluding the length of the tail fin). The length was then calculated by Matlab (R2021a), by converting pixel number to centimeters using the included reference ruler. All fish measured were siblings, and for blinding, genotypes were determined after the experiments.

### Fertility Analysis

Fish fertility was evaluated as described previously^12,14^. Briefly, 3-7 independent age-matched pairs of fish (one male, one female) of the indicated genotypes were placed in the same tank. All breeding pairs were allowed to continuously breed on sand trays, and embryos were collected and counted on a weekly basis. Results were expressed as the number of eggs per couple per week of egg-lay. Significance compared to the WT was calculated using repeated measures two-way ANOVA with a Sidak post-hoc.

### Histology

#### Hematoxylin and eosin

Tissues samples were processed as described previously^12–15,48–59^. Briefly, fish were euthanized with 500 mg/l tricaine (MS222, #A5040, Sigma). Paraffin sections were prepared by opening the body cavity of the fish and following a 72 h fixation in 4% PFA solution at 4°C, samples were dehydrated and embedded in paraffin using standard procedures. Sections of 5-10 μm were stained with Hematoxylin and Eosin, and examined by microscopy. A fully motorized Olympus IX23 microscope with an Olympus DP28 camera was used to collect images. Stages of egg and sperm development were identified as described previously^60^.

#### Immunohistochemistry

Fish were euthanized and dissected as previously described^13^. Muscle tissue was fixed in 4% PFA at 4°C for 2 hours, and immersed in an OCT-sucrose solution for 2 hours at 4°C. The OCT-sucrose solution is composed of 20% sucrose w/v (Bio-Lab #001922059100) and 30% OCT v/v (Scigen Scientific Gardena #4586) in PBS. Tissues were then transferred to OCT (12h at 4°C) and frozen in liquid nitrogen. All immersions were performed with mild shaking. Serial 10 µm sections were taken using a cryostat, airdried, and stored at -20°C. For immunostaining, slides were washed 3X in PBS, and permeabilized for 10 minutes in a permeabilization buffer containing 0.1% Triton (Avanator Performance materials #X198-07) and 1% BSA (Sigma-Aldritch #A7906 in PBS). Sections were then blocked for 10 minutes (DAKO #X0909), and incubation with primary antibodies overnight. The following primary antibody was used: rabbit anti-Nile Tilapia GH antibody (1:100), a generous gift from Prof. Berta Levavi-Sivan. After several washes, the sections were incubated for 1 h at room temperature with donkey anti rabbit Alexa Fluor 594 secondary antibody (Abcam #150064, 1:500) in antibody diluent (DAKO #S0809). After several washes, autofluorescence was quenched using TrueVIEW® autofluorescence quenching kit (Vector Labs #SP8500) according to the manufacturer’s protocol and mounted with VECTASHIELD® containing DAPI (Vector Labs #30326). Samples were imaged using a fully motorized Olympus IX23 microscope with a Photometrics BSI camera, and processed in imageJ^61^.

### Injection and ectopic over-expression of plasmids via electroporation

#### Cloning of killifish cDNAs

Killifish cDNAs were cloned from brain tissues from male and female killifish by homogenization in TRIreagent (Sigma #T9424) using 3 mm Nirosta disruption beads (PALBOREG FEDERAL #BL6693003000) and a tissue homogenizer (TissueLyser LT, Qiagen #85600). Total RNA was isolated from the lysed tissues using the Direct-zol RNA miniprep kit. (Zymo research #R2052), and the Verso cDNA Synthesis Kit (Thermo scientific #AB1453A) was used to prepare cDNA with random primers according to the manufacturer’s protocol. cDNA for the *gh1* and *fshb* was amplified using custom DNA oligonucleotides (Sigma) and Platinum SuperFi II DNA Polymerase (Invitrogen, #12361010). Primer sequences are available in Table S1. PCR products were purified (QIAquick PCR purification kit, Qiagen #28104) and sequence-verified. The sequence-verified ORFs were cloned using GIBSON (NEB, #E2611L) into the pLV-EGFP plasmid, which was modified such that each hormone is tagged with a GFP, separated by the T2A self-cleaving peptide ^20^. Plasmids and corresponding annotated maps are available via Addgene (## 194356, #194883).

#### In-vivo electroporation

The electroporation protocol was adapted from^16^. Fish were sedated in MS222 (200 mg/l), and 3-5 µl plasmid solution (Plasmid concentration 200-1000ng/µl)) containing Phenol Red for visualization (0.1% Sigma #P0290) was injected intramuscularly using a Nanofil syringe (WPI, #NANOFIL). Fish were then electroporated using 7 mm tweezer electrodes (NEP GENE #CUY650P10) with the ECM 830 generator. (BTX #45-0661). Electrodes were coated with wet cotton to minimize the risk to the fish. Fish were electroporated with 6 pulses of 28 V)14V for small fish, such as *gh1*^*Δ4/Δ4*^), for 60 ms each with an interval of 1 s between each pulse. Images were recorded by a Leica MC190HD camera mounted on a Leica M156FC microscope.

#### Doxycycline inducible expression

Fish were injected with a modified version of the <u>TLCV2</u> plasmid (Addgene #87360). This plasmid contains doxycycline induced Cas-9/GFP, which we modified to include Tol2 sites (Addgene #196331). Electroporation was performed as described above. Doxycycline treatment protocol was adapted from^62,63^. Briefly, Doxycycline hyclate (Dox, Sigma-Aldrich #D9891) was dissolved in 100% ethanol at 10 mg/mL for storage, and was added to fish water for a final concentration of 10 μg/mL. Fish were treated for 72 hours, and fresh water with Dox was changed every 24 hours. Due to possible light sensitivity of the drug, tanks were protected from light during treatment. Immediately following the treatment fish were imaged using a Leica MC190HD camera mounted on a Leica M156FC microscope.

